# The neurovascular coupling response of the aged brain is brain-state dependent

**DOI:** 10.1101/2024.07.09.602636

**Authors:** Xiao Zhang, Lechan Tao, Amalie H. Nygaard, Yiqiu Dong, Xiaoqi Hong, Carolyn M. Goddard, Chen He, Dmitry Postnov, Ilary Allodi, Martin J. Lauritzen, Changsi Cai

## Abstract

Brain aging lead to reduced cerebral blood flow and cognitive decline, but how normal aging affects neurovascular coupling (NVC) in the awake brain is unclear. Here, we investigated NVC in relation to calcium changes in vascular mural cells (VMCs) in awake adult and aged mice. We show that NVC responses are reduced and prolonged in the aged brain and that this is more pronounced at the capillary level than in arterioles. However, the overall NVC response, measured as the time integral of vasodilation, is the same in two age groups. In adult, but not in aged mice, the NVC response correlated with Ca2+ signaling in VMCs, while the overall Ca2+ kinetics were slower in aged than in adult mice. In particular, the rate of Ca2+ transport, and the Ca2+ sensitivity of VMCs were reduced in aged mice, explaining the reduced and prolonged vasodilation. Spontaneous locomotion was less frequent and reduced in aged mice as compared to young adult mice, and this was reflected in the ‘slow but prolonged’ NVC and vascular Ca2+ responses. Taken together, our data characterize the NVC in the aged awake brain as slow but prolonged, and underscoring the importance of brain state in understanding age-related mechanisms.

## Introduction

In the healthy adult brain, neurons rely on an on-demand system known as “neurovascular coupling” (NVC) to translate neural activity into a rapid increase in local blood flow, delivering blood-borne nutrients and expelling metabolic waste ^1^. It is acknowledged that age-related alteration in cerebral vascular function leads to a misalignment between increased metabolic needs and inadequate blood supply ^2–5^. Therefore, some studies suggest an impaired and declined NVC function in the aging brain. However, there are also compensatory neuronal strategies to counteract the age-associated alteration. Evidences from neuropathology and neuroimaging suggest the aging brain is not necessarily dysfunctional but remodeled, and can protect against function decline, by e.g. neural network reorganization, rise of neural activity efficiency and adaptive metabolic changes ^6–9^. Similarly, some studies have reported that NVC is not compromised in the aged brain and suggested the existence of adaptive adjustments ^10, 11^. The discrepancy may arise from different brain states in the experimental animals – neuronal and vascular activities are significantly different in the awake state compared to during sleep or under anesthesia ^12^. Anesthesia leads to profound decrease in baseline metabolism and affects hemodynamic responses ^13, 14^. Furthermore, it also affects young and old mice differently due to age-related alterations in body fat and liver function ^15, 16^. Therefore, employing awake animals to study age-related alterations in NVC to compare with anesthetized state is crucial, as a precise and comprehensive understanding of neurovascular coupling under a physiological condition will promote its translation into clinical studies.

Vascular mural cells (VMCs), which include smooth muscle cells at arteries, and pericytes at capillaries, play major roles in maintaining and regulating the vascular tone and cerebral blood flow ^17–19^. Specifically, the microvascular inflow tract (MIT), which comprises penetrating arterioles (PA) and arteriolar capillaries are the key functional compartments in regulating blood flow and maintaining hemodynamic homeostasis at capillary level ^4^. Further, pericytes at the MIT are contractile with unique sheathed or meshed morphology, which can be distinguished based on their locations and morphologies ^20–22^. Intracellular calcium level is essential in generation and regulation of myogenic tone in smooth muscle cells and contractile pericytes, mainly via a calcium-dependent process known as “excitation-contraction coupling”. Alteration of intracellular calcium activities in VMCs may contribute to the vascular aging in the central nervous system ^23,24^. Previous studies reported that in aged anesthetized mice, the vascular responsivity decreased at the first capillaries branching from arterioles, contributing to neurovascular uncoupling ^4^. However, whether this decline is associated with calcium signaling in contractile pericytes is still unclear. Therefore, it is necessary to re-examine in fully awake mice how, and to what extent, NVC is impaired in aged brains, and whether it is associated with alterations of calcium signaling in VMCs.

Here, using laser speckle contrast imaging and two-photon microscopy in adult and old awake behaving mice, we reported that in response to physiological and natural stimulations, NVC was preserved rather than impaired with age, with preserved total volume of functional hyperemia. Vascular responses in aged mice were reduced in amplitude but prolonged in time, which were functionally associated with slow but prolonged calcium dynamics and reduced calcium sensitivity in VMCs, and structurally with retracted coverage of contractile mural cells.

## Materials and Methods

### Animals

All procedures were approved by the Danish National Ethics Committee according to the guidelines set forth in the European Council’s Convention for the Protection of Vertebrate Animals used for Experimental and Other Scientific Purposes and are in compliance with the ARRIVE guidelines. All mice were kept on a C57BL/6 background and under a 12–12 hr light–dark cycle with food and water provided ad libitum from the cage lid. A total of 14 aged mice (20-24 month old) and 18 adult mice (3-4 month old) of both gender were used in this study. Acta2-GCaMP8.1/mVermilion (Tg(Acta2-GCaMP8.1/mVermilion)B34-4Mik, #032887) mouse strain were obtained from The Jackson Laboratory (USA).

### Chronic cranial window surgery

Chronic cranial window surgery was performed based on an established procedure ^25^. In brief, chronic window implantation was performed under aseptic conditions and isoflurane anesthesia. The mouse was placed in a stereotaxic frame on a heating pad (37°C). 5□mg/kg of Carprofen and 4.8□mg/g of dexamethasone with 0.5 mL saline were administered subcutaneously. Craniotomy was performed over the left somatosensory cortex (AP: -0.5, ML: -3; Ø□=□3□mm). The bone flap was carefully removed and the exposed brain temporarily covered with a haemostatic absorbable gelatine sponge prewetted with ice-chilled aCSF. The cranial opening was filled with aCSF, and then sealed using an autoclave sterilized round imaging coverslip (Ø□=□5□mm). A lightweight stainless steel head plate (Neurotar®) was positioned on the top of the skull surrounding the cranial window. The skull was coated with dental cement to secure the exposed bone was fully covered and the metal plate was firmly attached to the head. After the surgery, the animals were injected with 0.05□mg/kg of buprenorphine and 0.5 mL saline and returned to the cage on a pre-warmed heating pad. Postoperative care consisted of subcutaneous injections of carprofen and buprenorphine (2 and 4 days□post-surgery, dose as before). In the 7□days post-surgery, the convalescent animals were carefully monitored and their body weight were recorded every day.

### Laser speckle contrast imaging

Laser speckle contrast imaging was performed in adult and old awake mice with pre-implanted chronic cranial window. The recording hardware and software was customized as pervious describe ^26^. In brief, laser source was generated using a VHG-stabilized laser diode (LP785-SAV50, Thorlab) powered by a controller (CLD1011LP, Thorlab). Recording was performed using a CMOS camera (acA2040-90umNIR, Basler) and a ×5 0.15-NA objective (HCX PL FLUOTAR, Leica). Images were acquired with pixel resolution of 3.19 μm/pixel and image size of 1024*1024 at 50 frames per second. The awake mouse with chronic cranial window was head-fixed with free moving limbs. Two cameras were used to monitor facial behaviors (pupillometry, whisking) and body behaviors (locomotion, grooming). The real-time locomotion speed was recorded using a tracer placed beneath the air-floating cage. Air-puff stimulation was conducted to the distal end of mouse whiskers through a metal tube. A whisker air puff lasting 1 sec (3 Hz, 0.25 sec) was delivered after 30 sec baseline recording. Awake mouse pupil size, locomotion and whisking were recorded simultaneously with LSCI. The laser speckle images were post-processed using Matlab 2023a with custom-made code ^27^.

### Two-photon imaging

Before two-photon imaging, 50 μL of 4% wt/vol Texas Red-dextran (70,000 MW, Invitrogen) was applied intravenously to stain the vessel lumen. Fast repetitive hyper-stack imaging (4D imaging) was performed using a commercial two-photon microscope (Femto3D-RC microscope and Femtonics MES v6 software, Femtonics) equipped with a ×25 1.0-NA piezo motor objective. This method compensates for focus drift and allows for evaluation of the vasculature spanning a certain z-axis range. Each image stack was acquired within 1 sec and comprised 5–12 planes with a plane distance of 3–5 μm. This approach covered the whole z-axis range of the investigated blood vessels. The pixel sizes in the x–y plane were 0.2–0.38 μm. The excitation wavelength was set to 920 nm. The emitted light was filtered to collect green and red light from GCaMP8.1 (Acta2 positive mural cells) and Texas Red-dextran (vessel lumens), respectively. For awake mice imaging, whisker air-puff stimulation lasting 1 sec (3 Hz, 0.25 sec) was delivered after 30 or 40 sec baseline recording, while the camera monitored whether the stimulation elicited noticeable movement (locomotion speed > 1 cm/s or the obvious limb movement like grooming and lifting the limb) in mice. Only data where stimulation did not elicit noticeable movement were retained. Total recording time for each awake mouse in one experiment will not exceed more than 1.5 hr to avoid emotional harm. For anesthetized mice imaging, a mixture of ketamine (60 mg/kg) and xylazine (10 mg/kg) was given by i.p. injection to induce anesthesia with supplemental doses of ketamine (30 mg/kg) every 20 min.

### Immunohistochemistry and image analysis

Mice were anesthetized and intracardially perfused with a solution of normal saline for 3 min, followed by 4% paraformaldehyde (PFA) in 0.1 M PB for 2 min. Brains were removed and post-fixed in 4% PFA at 4 °C for 3 hr and then cut into 60-μm-thick coronal sections including primary somatosensory cortex. Slices were then incubated in permeabilizing buffer (0.3% Triton X-100 in PBS) for 15 min and blocked with donkey serum (S30-M, Sigma) (10% in PBS with 0.1% Triton X-100) for 2 hours at RT. All the sections were incubated overnights at 4 °C with primary antibodies including: rat anti podocalyxin (1:400, MAB1556, R&D Systems), goat anti PDGFRβ (1:250, AF1042, R&D Systems), mouse anti αSMA-FITC (1:200, F3777, MilliporeSigma). The sections were then washed in 0.1% PBST (0.1% Triton X-100 in PBS) for 15 min three times and incubated at room temperature for 2 hours with corresponding secondary antibodies including: donkey anti goat 568 (1:500, A-11057, Invitrogen), donkey anti rat 647 (1:500, A78947, Invitrogen). Hoechst 33342 (1:4000 dilution) was applied together with secondary antibodies. After washing with 0.1% PBST for 15 min three times, the sections were mounted on glass slides in SlowFade Diamond Antifade Mountant (Invitrogen, S36963) and cover glasses were sealed with nail polish. All images were acquired on a Zeiss LSM 900 confocal microscope equipped with Plan-Apochromat ×20/0.8 DRY objective at the Core Facility for Integrated Microscopy, University of Copenhagen. Image analysis was performed by FiJi (US National Institutes of Health). Different channels were thresholded and cell numbers were determined according to the Hoechst 33342 channel.

### Computational modeling

We assume that the diameter data follow roughly a Gaussian function, and the calcium data follow the derivative of a Gaussian. Therefore, we use the functions defined as

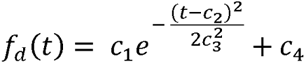

and

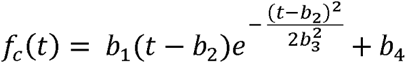

to fit the diameter data and the calcium data in the least-squares sense, respectively. In the formulas, *c*_1_ and *b*_1_ are the amplifiers, *c*_2_ and *b*_2_ represent the mean, *c*_3_ and *b*_3_ are positive and denote the standard variation, and c_4_ and b_4_ are the bias.

After finding the fit functions, we try to discover the relation between the calcium data and the diameter data. We assume that the diameter fit function f_d_ (t) and the calcium fit function f_c_(t) satisfy the relation:

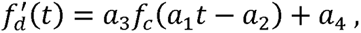

where *a*_1_ and *a*_2_ give the time transform and denote the acceleration and the shift, respectively. Further, a_3_ shows the scaling difference and a_4_ gives the bias difference. According to the formulas of *f_d_*(*t*) and *f_c_*(*t*), we can form their relation through

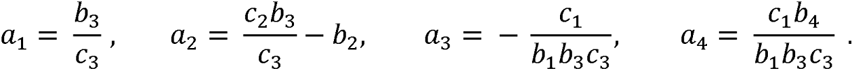

### Pupillometry and whisking detection

The video of mouse face at the right side covering pupil and whiskers was recorded by a CMOS camera (GO-5100M-PGE, JAI), with frame rate of 30 Hz and frame size of 800 * 800 pixels. Both the pupillometry and whisking movement were analyzed offline. Pupillometry was measured under a constant light and analyzed by DeepLabCut, assigning eight dots to define the contour of a pupil. The size of a pupil was calculated by the area delineated by the eight dots at each frame. Whisking was analyzed by customized MATLAB code. The strength of cross-correlation was calculated between two consecutive frames, representing the movement intensity of whisking. This algorithm precisely detected the on and off time of whisking movement. Two consecutive whisking events with inter-event-interval less than 0.25 sec were merged as single event.

### Quantification and statistical analysis

All experiments were conducted independently with a minimum of three biological replicates, as specified in the figure legends. Bar and line graphs were generated using Origin 2020 and compiled using Adobe Illustrator 2022. GraphPad InStat 3 was used for the formal tests for normality to assess data distribution and further statistical analyses. For the comparison of two groups, we used either a student unpaired 2-tailed t-test with or without Welch correction, or Mann-Whitney nonparametric test, depending on whether the data follows a normal distribution and the populations have equal SDs. Statistical significance was set at *p < 0.05, **p < 0.01 and ***p < 0.001. ‘n.s’ indicates no significant statistical difference (p > 0.05). All quantitative data are presented as mean ± SD.

## Results

### Reduced but prolonged blood flow responses upon whisker air puff in old awake mice

To examine the regional neurovascular coupling function in adult and old mice, we used laser speckle contrast imaging (LSCI) to measure the cortical blood flow changes during whisker air-puff stimulation in awake mice ^28^. After adaptive training, mice were head-fixed and mounted on the imaging device via a head bar, with freely moving limbs on an air-cushioned mobile cage. Whisker air-puff stimulation was administered to the distal end of the mouse whiskers through a puffing tube (Figure 1a and Figure S1A). In both adult (3-4 month old) and old (> 20 month old) mice groups, whisker air-puff stimulation induced pupil dilation of similar amplitude and duration (Figure S1B-E). In response to stimulation, blood flow increased in the primary somatosensory cortex and adjacent regions of both young adult and aged mice. The response zone was defined as a pixel intensity increase of at least 5% relative to the baseline after stimulation (Figure 1b-d). The maximum area and spread speed of the response zone were measured and results showed no significant differences between the two age groups (Figure S2A-E). In old mice, the peak amplitude of the cortical blood flow rise in the response zone was decreased compared to young adult mice (Adult: 19.84% ± 4.89% v.s Old: 14.76% ± 3.63%, **p < 0.01), and the half-peak duration of the blood flow rise was longer (Adult: 4.95 ± 0.61 sec v.s Old: 6.45 ± 2.23 sec, *p < 0.05) (Figure 1e-g). There was no significant difference between the two age groups in half peak latency and the area under the curve (AUC, i.e. time integral of vasodilation) (Figure 1h and 1i), which reflect the response onset time and total volume of blood flow change, respectively. This indicates that the total volume of the functional hyperemia was preserved while the duration was prolonged with age.

**Figure 1.**
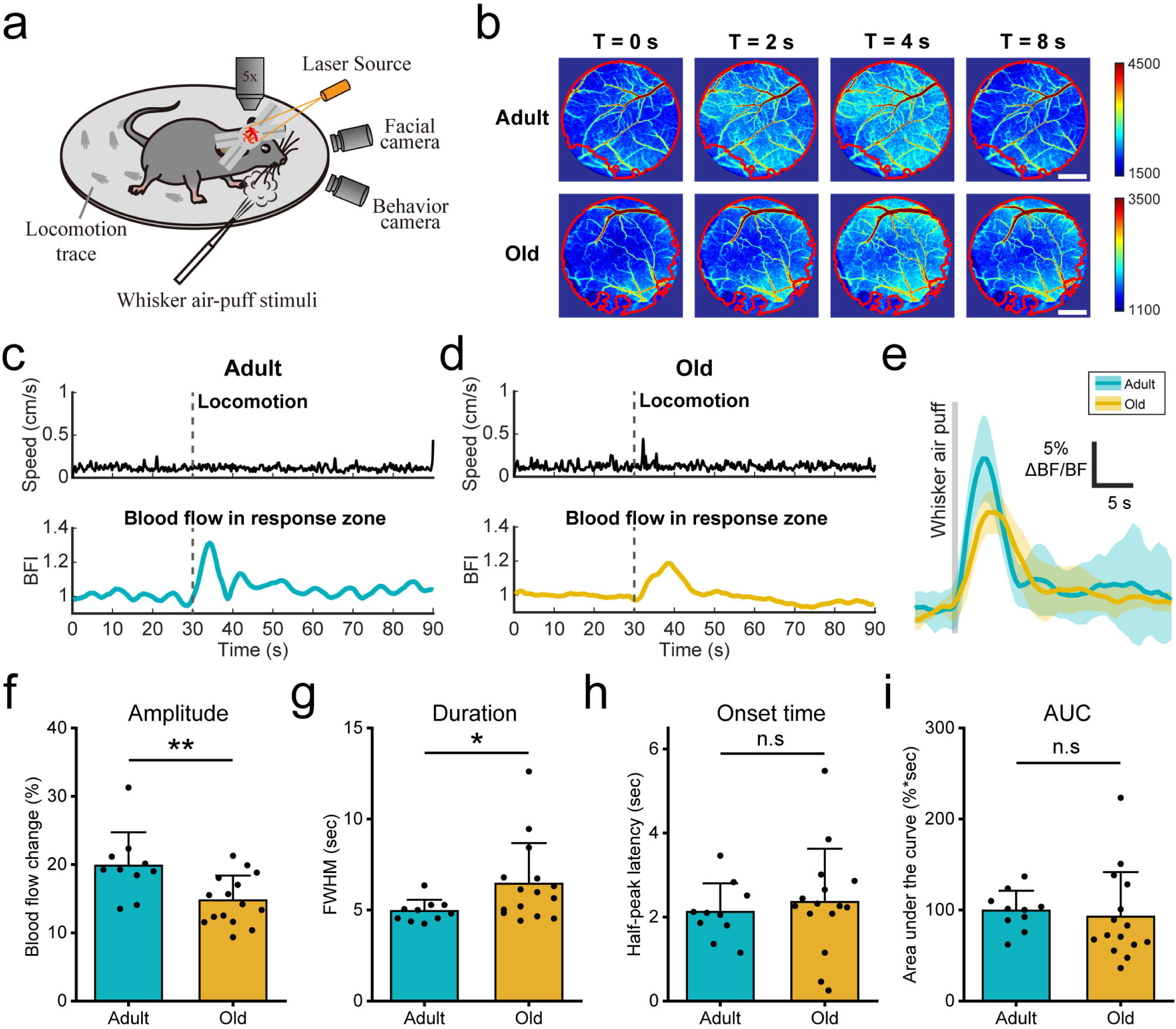
Reduced but prolonged functional hyperemia in the aged brain. (a). Schematic for laser speckle contrast imaging set-up with awake mice. (b). Representative images of cortical blood flow change in response to air-puff stimulation in adult and old mice. The response zone is marked with a red circle. Scale bar, 1 mm. (c and d). Representative traces of locomotion speed and blood flow index upon air-puff stimulation in adult and old mice. The dashed line indicates the time when the air-puff was delivered. (e). Mean (solid curve) and SD (shadow) traces of the blood flow change upon air-puff stimulation in adult and old mice. (f-i). Comparison of amplitude (f), duration (g), onset time (h) and area under the curve (i) of blood flow index change upon air-puff stimulation in adult and old mice. N= 10 recordings from 7 adult mice and 15 recordings from 7 old mice. Data are presented as mean values ± SD. In (f, h and i), unpaired t test; in (g), Mann-Whitney test. *p < 0.05; **p < 0.01; n.s, not significant.

To identify the vascular elements contributing to the changes of NVC in aged mice, we placed multiple region of interests (ROIs) at two different types of segments in the LSCI cortical images, i.e. the pial artery region and the parenchyma region (mainly consisting of capillaries) (Figure 2a and 2g). Changes of pial vessel diameter and intraluminal blood flow at the pial artery/parenchyma regions were analyzed for each ROI. Whisker air-puff stimulation induced similar changes in dilation amplitude, half-peak duration, onset time, and AUC of pial artery dilation in adult and old mice (Figure 2b-f). However, compared with young adult mice, stimulation in old mice revealed decreased amplitude and prolonged duration of blood flow changes in the parenchymal region but not in the pial artery region (Figure 2h-j), with no change in AUC and onset time of blood flow response (Figure 2k and 2l). This suggests that alterations in capillary mechanisms may be the crucial for neurovascular coupling remodeling in old mice.

**Figure 2.**
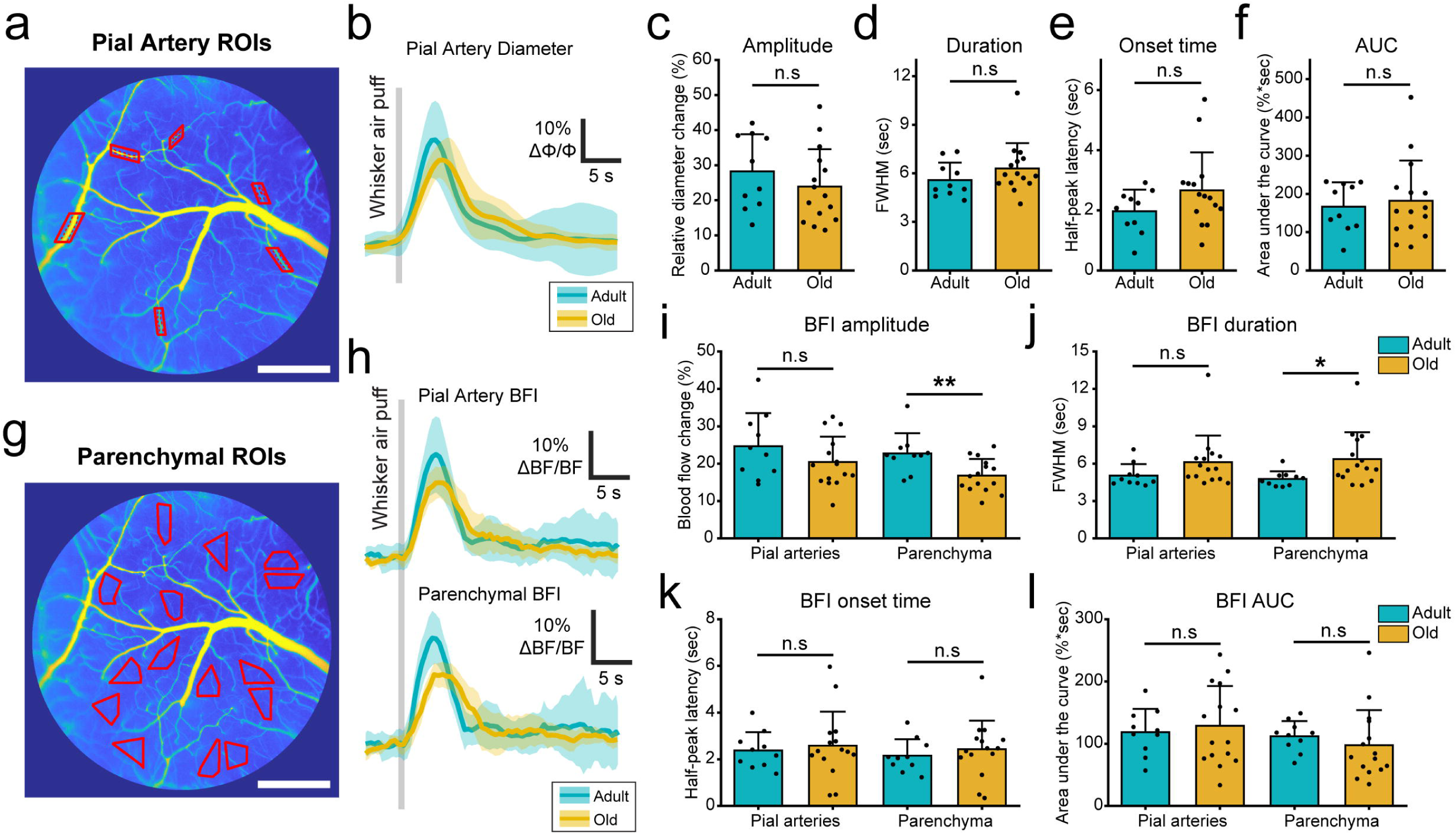
Age-related changes in functional hyperemia mainly occur in brain parenchyma. (a). The representative image illustrating the selection of pial arteries in LSCI. Six ROIs were randomly placed on pial arterial locations within the responsive zone, ensuring minimal interference from venous blood flow within the ROIs. Scale bar, 1 mm. (b). Mean (solid curve) and SD (shadow) traces of the pial artery diameter change upon air-puff stimulation in adult and old mice. (c-f). Comparison of dilation amplitude (c), duration (d), onset time (e) and area under the curve (f) of pial artery diameter change upon air-puff stimulation in adult and old mice. N= 10 recordings from 7 adult mice and 15 recordings from 7 old mice. Data are presented as mean values ± SD. (g). Representative image showing the selection of cortical parenchyma regions in LSCI. Scale bar, 1 mm. (h). Mean (solid curve) and SD (shadow) traces showing the blood flow change in both pial artery region and parenchymal region in response to air-puff stimulation in adult and old mice. (i-l). Summary graphs showing the amplitude (i), duration (j), onset time (k) and area under the curve (l) of blood flow change in both pial artery region and parenchymal region upon air-puff stimulation in adult and old mice. N= 10 recordings from 7 adult mice and 15 recordings from 7 old mice. Data are presented as mean values ± SD. In (c, d, f, i, j _right_, k and l), unpaired t test; in (e, j _left_), Mann-Whitney test. *p < 0.05; **p < 0.01; n.s, not significant.

Next, we used *in vivo* two-photon microscopy to examine the neurovascular coupling function at the first few branching capillaries from penetrating arterioles ^29^. We classified the capillaries by their branching orders, with the first order being the first capillary branching from the penetrating arterioles (PA), and so forth (Figure 3a). We utilized the Acta2-GCaMP8.1/mVermilion transgenic mice expressing calcium fluorescent indicator GCaMP8.1 in contractile mural cells and intravenously injected Texas Red dextran to label the vessel lumen and identified the MIT (Figure 3a). At resting state, the basal vessel diameter of awake old mice showed no significant difference compared to adult mice (Figure S3A). Upon whisker air-puff, vasodilation occurred across the vasculature of the MIT within the somatosensory cortex (Figure 3b and 3c). PA in the old brains showed a similar level of vasodilation compared to adult mice. However, a decreased dilation amplitude was found at the capillaries level in old mice, especially at the first order capillary (1st Cap), where the dilation was ∼26% smaller as compared with young adult mice (Adult: 32.45% ± 11.41% v.s Old: 24.01% ± 11.92%, *p < 0.05) (Figure 3d). Further, the dilation duration and half peak latency at capillaries but not at PA were prolonged in old mice, while the AUCs remained unaltered across all the vasculature (Figure 3e-g), indicating capillaries are the key vascular location with altered functional hyperemia profile in the aged brain. We further calculated the rate of change of vasodilation and vasoconstriction and found that in old mice, the rate of dilation was decreased by ∼22%, ∼26% and ∼36% and the return to normal was decreased by ∼26%, ∼35% and ∼29% at PA, 1st Cap and 2nd Cap respectively as compared to young adult mice (Figure S3B and S3C). This suggested a slower speed of vasodilation and vasoconstriction across all the vasculature in MIT in old mice.

**Figure 3.**
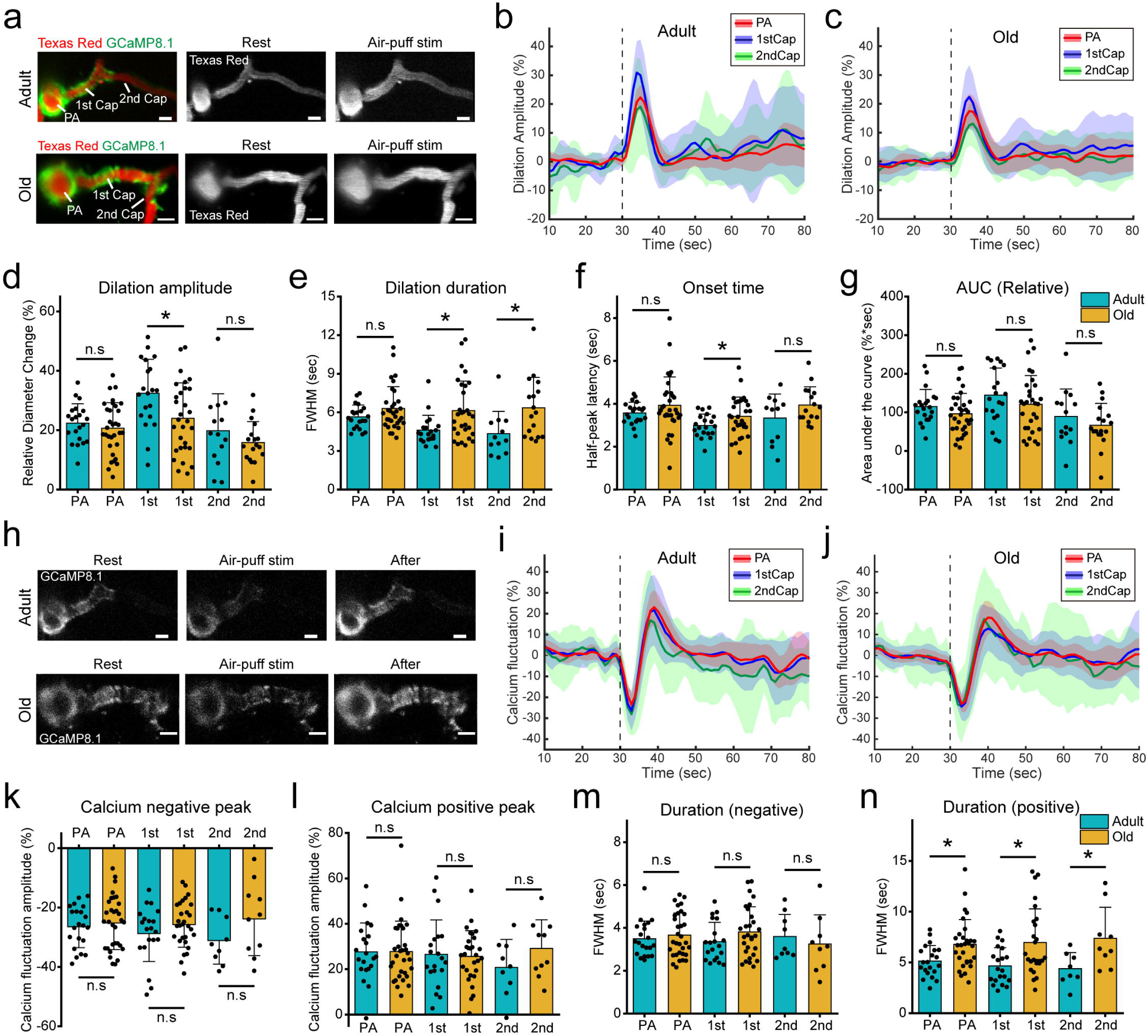
Stimulation evoked vasodilation and vascular mural cell calcium dynamics are prolonged in aged MIT. (a). Representative images of MIT, which contain penetrating arterioles (PA), first and second order capillaries (1st Cap & 2nd Cap), in response to air-puff stimulation in adult and old Acta2-GCaMP8.1 mice. The vessel lumen was labelled with Texas Red dextran, and the vascular mural cell expressed GCaMP8.1. Scale bar, 10 μm. (b and c). Mean (solid curve) and SD (shadow) traces of the vessel diameter change at each vasculature location upon air-puff stimulation in adult and old mice. The dashed line indicates the time when the air-puff was delivered. (d-g). Comparison of relative dilation amplitude (d), dilation duration (e), onset time (f) and area under the curve (g) at each location upon air-puff stimulation in adult and old mice. N = 21 vessels from 8 adult mice and 32 vessels from 7 old mice. Data are presented as mean values ± SD. In (d _PA & 1st_, e _2nd_ and g), unpaired t test; in (d _2nd_ and f _PA & 1st_), unpaired t test with Welch correction; in (e _PA & 1st_ and f _2nd_), Mann-Whitney test. *p < 0.05; n.s, not significant. (h). Representative images of calcium fluctuation in vascular mural cell at each vessel location in response to air-puff stimulation in adult and old Acta2-GCaMP8.1 mice. Scale bar, 10 μm. (i and j). Mean (solid curve) and SD (shadow) traces of the calcium fluctuation in VMCs at each vasculature location upon air-puff stimulation in adult and old mice. The dashed line indicates the time when the air-puff was delivered. (k-n). Bar graphs summarizing the amplitude and duration of both negative and positive peaks of calcium fluctuation in VMCs upon air-puff stimulation in adult and old mice. N = 21 vessels from 8 adult mice and 32 vessels from 7 old mice. Data are presented as mean values ± SD. In (k, l, m 2nd and n 2nd), unpaired t test; in (m _1st_), unpaired t test with Welch correction; in (m _PA_ and n _PA & 1st_), Mann-Whitney test. *p < 0.05; n.s, not significant.

### Preserved amplitude and prolonged time course of calcium signals in aged vascular mural cells

Calcium plays a critical role in the regulation of contraction and relaxation of smooth muscle cells and contractile pericytes. The calcium dynamics in VMCs were measured in the Acta2-GCaMP8.1 mice. In both adult and old mice, whisker air-puff stimulation elicited a biphasic calcium response in VMCs, which started with a rapid [Ca^2+^]_i_ drop and followed by a [Ca^2+^]_i_ rise (Figure 3h-j). In contrast to the reduced and prolonged vessel diameter change at capillaries, the negative and positive peak amplitudes of the calcium signals was the similar in adult and old mice (Figure 3k and 3l). However, the vascular mural cell calcium response lasted longer in old than in young mice, suggesting a slower calcium dynamic in the aged brain (Figure 3m and 3n). We further compared the rate of calcium changes in VMCs in the two age groups and found it was significantly decreased at both PA and 1st Cap in aged mice (Figure S3D-H), indicating slower calcium dynamics with age.

Further, to compare the difference of vascular reactivity between awake and anesthetized states, we measured the vessel diameter change and calcium fluctuation in VMCs in response to whisker-pad stimulation in anesthetized Acta2-GCaMP8.1 mice (Figure S4A-D). Consistent with our previous research findings ^4^, vessel dilation amplitude and AUCs at both PA and capillary levels declined with age (Figure S4E and S4F). Unlike the observation in awake animals, the response of vascular mural cell calcium signal to whisker stimulation was significantly reduced in old mice compared to adult mice under anesthesia (Figure S4G and S4H), indicating that anesthesia has major effects on calcium-dependent vascular mural cell responsivity. This indicates that the effect of aging on calcium dynamics in VMCs is different between awake and anesthetized brain states.

### Spontaneous locomotion and physiological whisking induced unique calcium-dependent vascular responses

While whisker air puff which may elicit a startle effect, spontaneous locomotion of awake mice is a natural and physiological behavior, which induces neurovascular coupling ^25^. Further, locomotion is also a vulnerable factor with regard to aging ^30^. To examine whether spontaneous locomotion elicits different neurovascular coupling in adult and old mice, we compared the vascular responsivity and calcium dynamics between the two age groups (Figure 4a-c). The old mice exhibited a reduced intensity of spontaneous locomotion, with a decrease of the average speed and travel distance, and a similar motion duration for each locomotion event (Figure 4d-f). Unlike air-puff stimulation, the neurovascular coupling elicited by spontaneous locomotion was significantly weakened in old mice as indicated by a decrease in the dilation amplitude across all the vasculature of the MIT and reduced AUC of vessel dilation in PA (Figure 4g and Figure S5A-C). Interestingly, although the vasodilation duration did not show a significant difference between the two age groups, correlation analysis results suggested that the same duration of locomotion led to longer vasodilation in old mice (Figure 4h and Figure S5D). This suggests a vascular function remodeling in the old brains, which is characterized by a reduced but prolonged vasodilation. Vascular mural cell calcium dynamics accompanying spontaneous locomotion was also compared in the two age groups (Figure 4i-k), but showed no differences between adult and old mice (Figure 4l-o). However, old mice still exhibit a longer duration of vascular mural cell calcium signals in response to same spontaneous locomotion time (Figure S5E).

**Figure 4.**
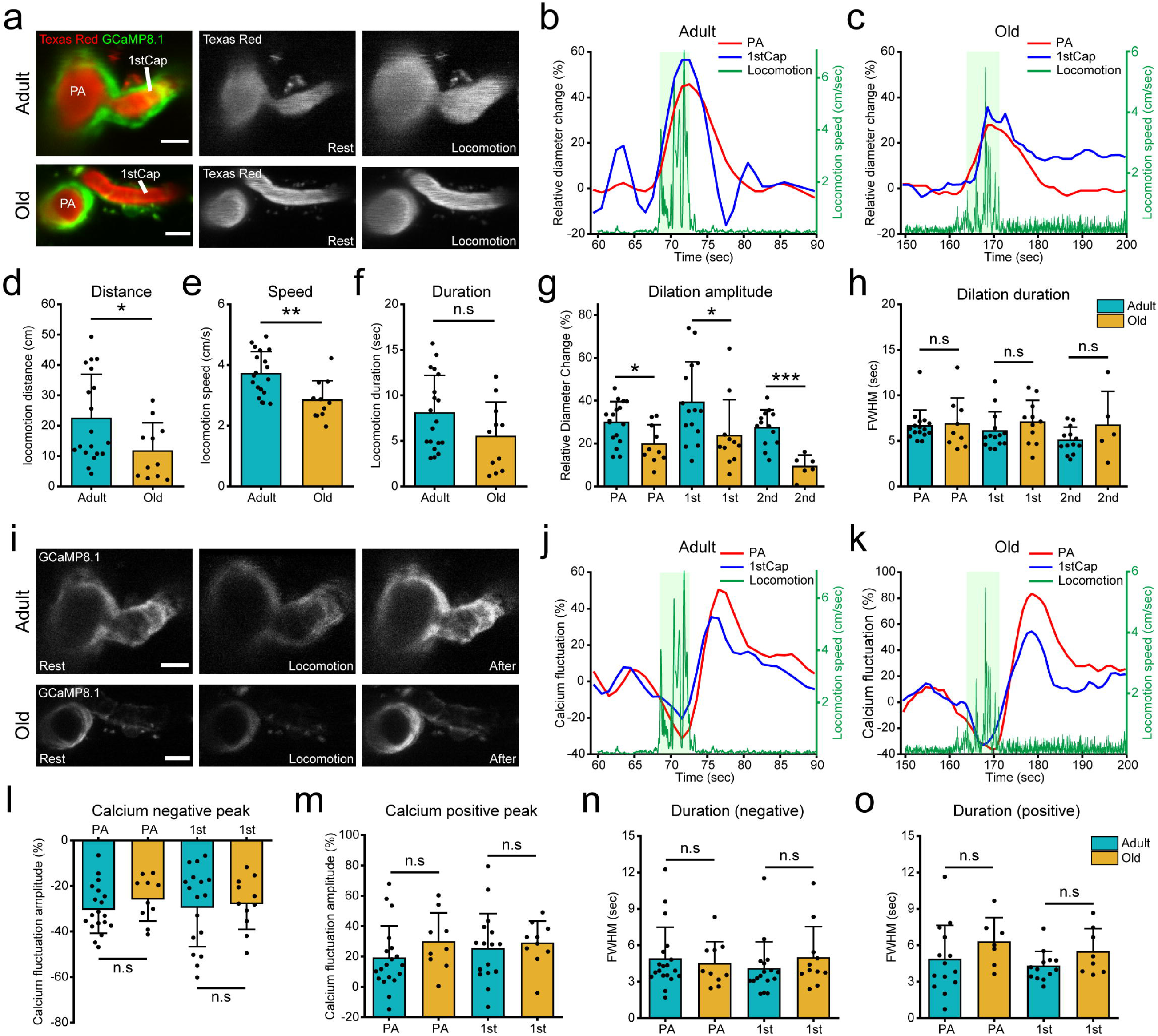
Locomotion induces a reduced vasodilation but preserved mural cell calcium dynamics in the aged brain. (a). Representative images of spontaneous locomotion induced vessel dilation at penetrating arterioles (PA) and first order capillaries (1st Cap) in adult and old Acta2-GCaMP8.1 mice. Note the vessel lumen was labelled with Texas Red dextran, and the vascular mural cell expressed GCaMP8.1. Scale bar, 10 μm. (b and c). Representative traces showing vessel dilation response to spontaneous locomotion. The green shadow indicates that the mouse is in spontaneous locomotion. Note that one spontaneous locomotion is defined as starting from when the locomotion speed exceeds 1 cm/s until the end of the continuous movement. (d-f). Summary graphs showing the locomotion parameters including average distance, speed and duration from single movement in adult and old mice. N = 19 recordings from 6 adult mice and 11 recordings from 4 old mice. Data are presented as mean values ± SD. In (d and f), Mann-Whitney test; in (e), unpaired t test. *p < 0.05; **p < 0.01; n.s, not significant. (g and h). Comparison of relative dilation amplitude (g) and dilation duration (h) of spontaneous locomotion induced vessel dilation in adult and old mice. N = 19 vessels from 6 adult mice and 11 vessels from 4 old mice. Data are presented as mean values ± SD. In (g _PA & 2nd_), unpaired t test; in (h _2nd_), unpaired t test with Welch correction; in (g _1st_ and h _PA & 1st_), Mann-Whitney test. *p < 0.05; **p < 0.01; ***p < 0.001; n.s, not significant. (i). Representative images of spontaneous locomotion induced calcium fluctuation in vascular mural cell at each vessel location in adult and old Acta2-GCaMP8.1 mice. Scale bar, 10 μm. (j and k). Representative traces showing calcium signal fluctuation in VMCs in response to spontaneous locomotion. The green shadow indicates that the mouse is in spontaneous locomotion. (l-o). Bar graphs showing the amplitude and duration of negative and positive peak of calcium fluctuation which induced by spontaneous locomotion in VMCs in adult and old mice. N = 19 vessels from 6 adult mice and 11 vessels from 4 old mice. Data are presented as mean values ± SD. In (l, m and o), unpaired t test; in (n), Mann-Whitney test. n.s, not significant.

Whiskers are the precise tactile terminals for mice to perceive the environment ^31^. We compared the whisking behavior-induced neurovascular coupling responses in adult and old mice at three different states: air-puff induced whisking, spontaneous whisking in the resting state, and whisking in association with spontaneous locomotion (Figure S6A and S6B). Whisker air-puff stimuli elicited whisking events of similar duration in the two groups (Figure S6C), while spontaneous locomotion was associated with shorter whisker movements in old than in adult mice (Figure S6D). In the resting state, old mice showed a slightly higher spontaneous whisking occurrence (∼37% of total time) than adult mice (∼30% of total time) (Figure S6E). We divided the whisking events at resting state into two distinct patterns, characterized as short-term (<2 sec) and long-duration (≥ 2 sec) whisking (Figure S7A and S7B). Short-term whisking did not induce any significant changes in blood flow and mural cell calcium, whereas long-duration whisking led to significant but similar blood vessel dilation and vascular mural cell calcium change in both adult and old mice (Figure S7C and S7D). These data suggested that whisker perception and related sensory input function was preserved in old mice.

### Calcium sensitivity in aged vascular mural cells is decreased at capillaries

Calcium regulation within VMCs is a fundamental process that governs their contractility. The sensitivity to the intracellular calcium changes in VMCs determines the speed and amplitude of vessel diameter change. To compare the calcium sensitivity of VMCs in adult and old mice, we performed linear regressions by correlating the AUC of negative and positive phases of vascular mural cell calcium dynamics with the AUC of rising and falling phases of vessel diameter changes. For vasodilation, there was a strong positive correlation between the negative calcium peak and vessel diameter change at PA and 1st Cap in adult mice (Figure 5a-c). However, old mice lost this calcium-diameter relationship at PA level and the correlation was weak for 1st Cap (Figure 5a-c). For vasoconstriction, adult mice showed a significant correlation between calcium diameter reduction for both PA and capillaries levels, whereas the old mice did not (Figure 5d-f). To further assess the age-related alteration of the relationship between vascular mural cell calcium and the vessel diameter change, we developed a simple mathematical model based on the physiological calcium and diameter waveforms upon whisker air puff obtained by *in vivo* two-photon imaging. The model assumed that the rate of change of vessel diameter was produced by the concomitant change in mural cell calcium (Figure 5g). Both diameter recordings and calcium were simulated based on Gaussian functions (Figure 5h and 5i), and the relationship of the two Gaussian functions were fitted with nonlinear regression. We considered the calcium function as input and diameter function as output and calculated the linear coefficients a1, a2, a3 and a4 in adult and old mice, representing the elongation of time, onset time delay and amplitude magnification and baseline drift, respectively for adult and old mice at PA, 1st Cap and 2nd Cap (Figure 5j-l). The results suggested that at different vascular locations and with the same calcium changes, the output diameter change of old mice can be characterized as: the duration of diameter change is prolonged (Figure 5j), the delay of diameter change from onset of calcium input is prolonged (Figure 5k), and the amplitude of diameter change is smaller, i.e. calcium sensitivity decreases (Figure 5l). These results are consistent with the outcomes of our two-photon experiments, and highlight the features of calcium-diameter correlation in VMCs of old awake mice, i.e. prolonged in time, and decreased in amplitude. Taken together, those results suggest calcium sensitivity of VMCs in old awake mice is reduced and may lead to a decrease in the vascular regulation efficiency.

**Figure 5.**
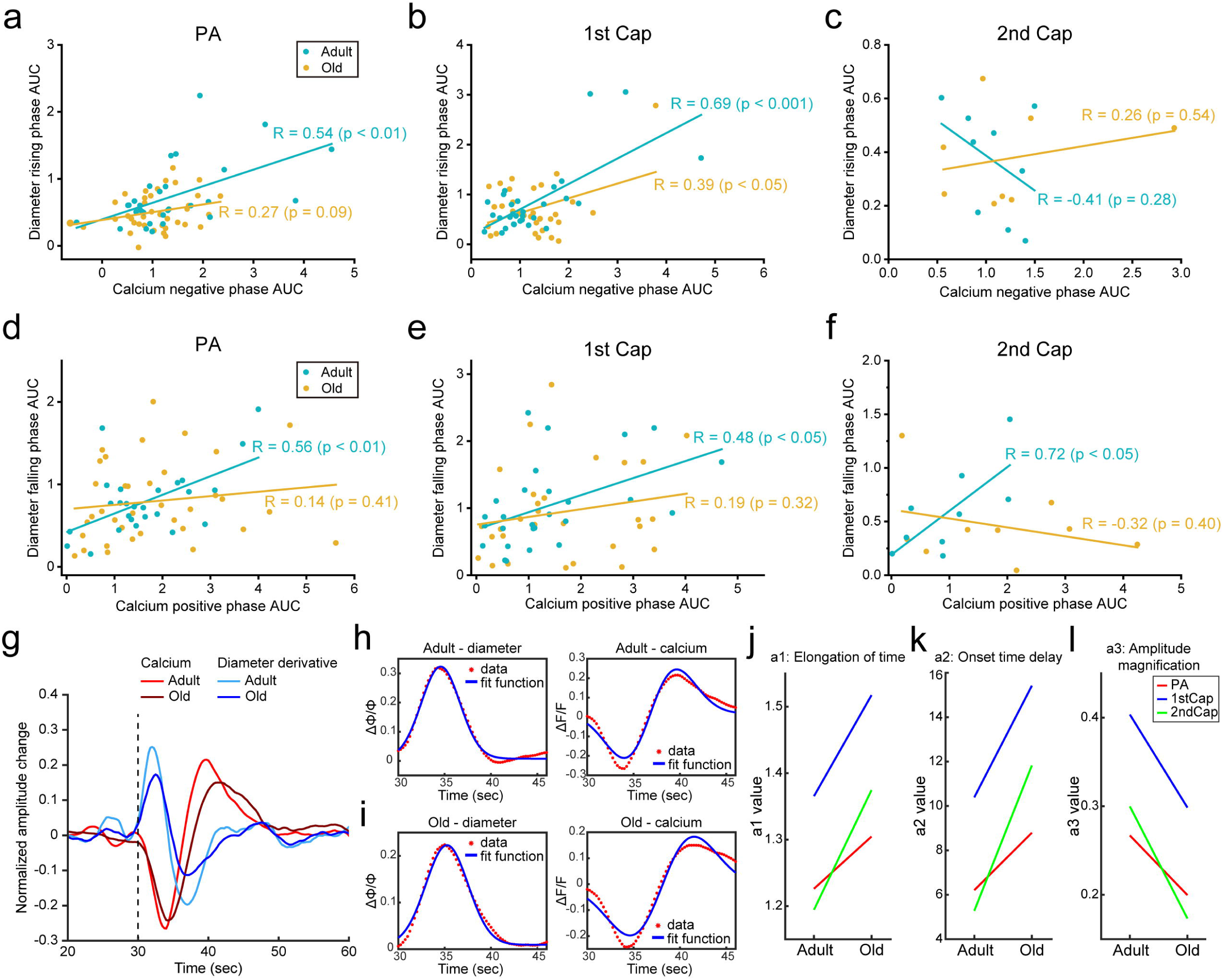
Vascular mural cell calcium sensitivity decreases in the aged brain. (a-c). Scatter plots summarizing the relationship between the area under the curve of mural cell calcium fluctuation negative peaks and the corresponding vessel diameter changes during vessel dilation phase (from baseline to peak). The correlation analysis is presented using linear fitting and Pearson’s r value is indicated. N = 30 vessels from 9 adult mice and 40 vessels from 7 old mice. (d-f). Scatter plots showing the relationship between the area under the curve of mural cell calcium fluctuation positive peaks and the corresponding vessel diameter changes during vessel constriction phase (from peak back to normal). The correlation analysis is presented using linear fitting and Pearson’s r value is indicated. N = 30 vessels from 9 adult mice and 37 vessels from 7 old mice. (g). Mean traces of the derivative of the diameter change (blue) and the calcium fluctuation in mural cell (red) upon air-puff stimulation in adult and old mice. (h and i). The linear relationship between the fit curve that simulated as Gaussian functions with the average vessel dilation curve and mural cell calcium fluctuation curve in adult and old mice. (j-l). Comparison of elongation of time (a1, j), onset time delay (a2, k) and amplitude magnification (a3, l) at each vessel location in adult and old mice with calcium function as input and diameter function as output. Note that a4 is not shown, because all data are normalized and baseline drift is minimal.

### Retraction of αSMA transition in old mice and decreased vascular responsivity

During aging, the cerebral vascular system undergoes both functional and morphological alteration ^32^. To further explain the decreased calcium sensitivity with age, one possibility is that a decrease in the coverage of contractile mural cells may lead to a reduction in vessel responsivity and regulatory efficiency. For this purpose, we examined the coverage of αSMA at the MIT to check whether it undergoes retraction in old mice. αSMA expression was observed along arteriolar capillaries, extending to the transition point where αSMA positive contractile mural cells transitioned both shape and function into αSMA negative non-contractile pericytes (Figure 6a). Using immunohistochemical staining of αSMA and endothelial cells, we measured the distributions and vessel diameters of these αSMA transition points across different order of capillaries in old mice as compared with adult mice. In old mouse brains, the location of the transition points shifted towards lower-order capillaries (Figure 6b and 6c) while the distance to the upstream PA decreased (Adult: 135.13 ± 73.92 μm v.s Old: 89.52 ± 57.41 μm, ***p < 0.001, Figure 6d), and the vessel diameters at the transition points increased with age (Adult: 4.74 ± 1.24 μm v.s Old: 5.25 ± 1.36 μm, ***p < 0.001, Figure 6e). This suggests that reduced coverage by contractile pericytes is found predominantly at capillary levels in old mice. Capillary pericytes play a crucial role in regulating blood flow during rises in brain activity and loss of pericytes could lead to a delayed vascular response upon neural stimulation ^33^, therefore the pericyte density was measured by counting PDGFRβ positive cell number on capillaries (Figure S8F). In line with the previous reports ^4^, the density of pericytes on blood vessels remained the same in young adult and old mice. However, the spatial density of pericytes decreased by approximately 20% in aged brains (Figure S8G and S8H). Taken together, these data suggested a retraction in the coverage of contractile VMCs on capillaries and a reduction of pericytes in old mice, which may partially explain the decreased responsivity of capillaries in old mice.

**Figure 6.**
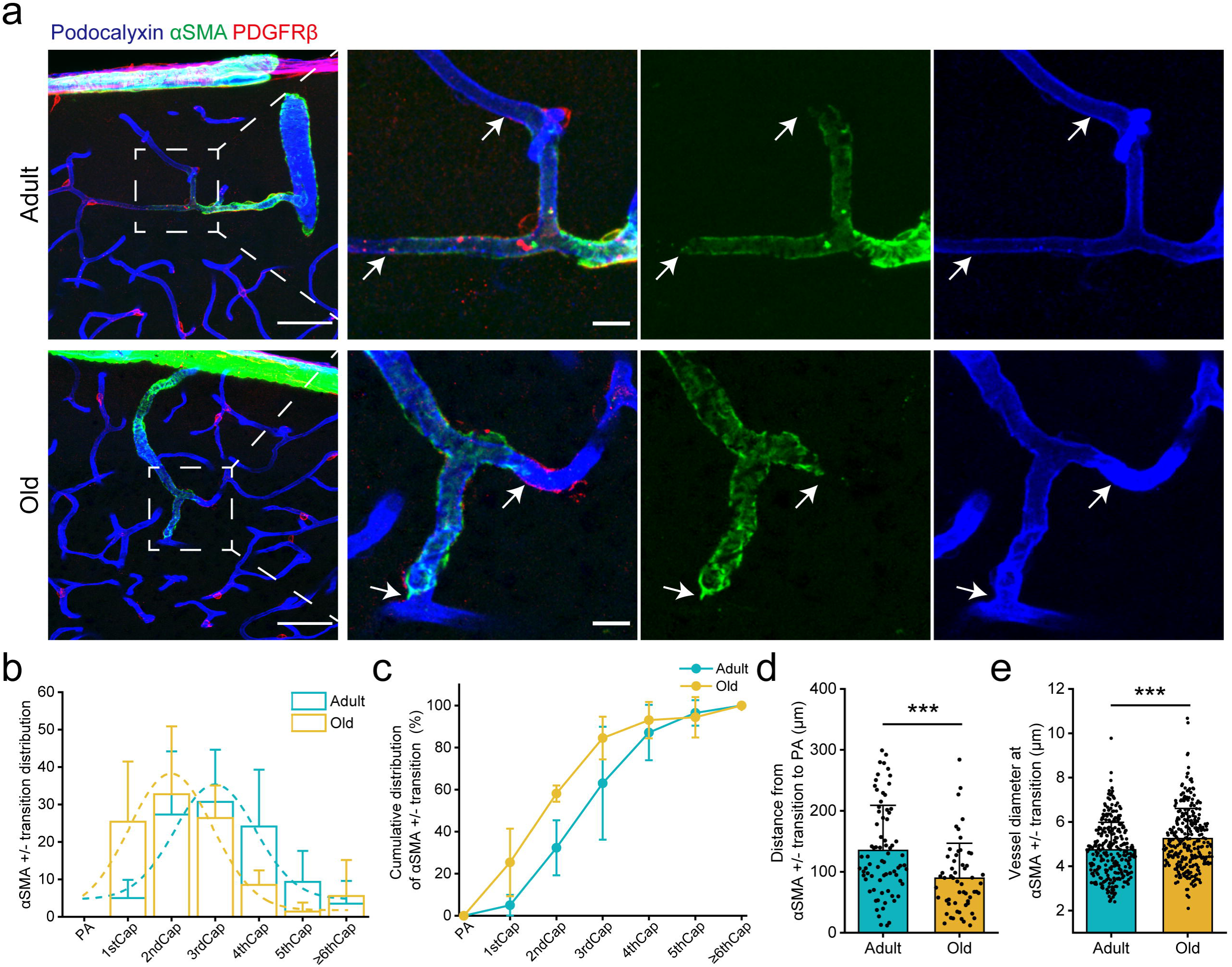
Capillary aSMA coverage undergoes retraction during aging. (a). Immunostaining of vascular mural cell in adult and old mice cortex. The arrows show the terminal positions of aSMA staining along the arteriolar capillaries. Scale bar: 50 μm (left) and 10 μm (inlet). (b and c). Histogram and cumulative distribution graph showing the probabilities of aSMA transition points distributed across each blood vessel locations in adult and old mice. The dashed lines are presented using nonlinear curve fitting (Gauss). N = 75 vessels from 3 adult mice and 57 vessels from 3 old mice. (d and e). Bar graphs summarizing the distance from the aSMA transition point to its upstream PA (adult: n = 75 vessels; old: n = 57 vessels) and the vessel diameter at each aSMA transition point (adult: n = 289 vessels; old n = 289 vessels) in adult and old mice. Data were collected from 3 adult and old mice and are presented as mean values ± SD. in (d and e), Mann-Whitney test. ***p < 0.001.

## Discussion

In this study, we reported an age-related remodeling of neurovascular coupling function in awake mice. In the aged mouse brain, vessel dilation is reduced in amplitude but prolonged in time, resulting in similar volume of functional hyperemia as in adult brains in response to a physiological stimulus. Using two-photon calcium imaging, we identified a decrease in calcium handling speed of VMCs, contributing to prolonged vascular dilation. This indicates that the mechanism underlying the age-related shift in NVC is associated with altered calcium kinetics in contractile VMCs. Furthermore, correlation analysis and mathematical modeling results suggested an altered relationship between mural cells calcium dynamics and vessel diameter change in the aged brain, accompanied by a decrease in calcium sensitivity in aged VMCs.

Moreover, we observed retraction of αSMA coverage on capillaries, implying a reduced contractile capability at the capillary level. Taken together, our results suggested that neurovascular coupling in the aged brain compensates for reduced vascular mural cell calcium sensitivity by slowing down the calcium transport speed, thereby extending the duration of vascular dilation and eventually reaching the same level of hyperemia. This remodeled mechanism may underlie the preserved NVC response in aged mice (Figure 7).

**Figure 7.**
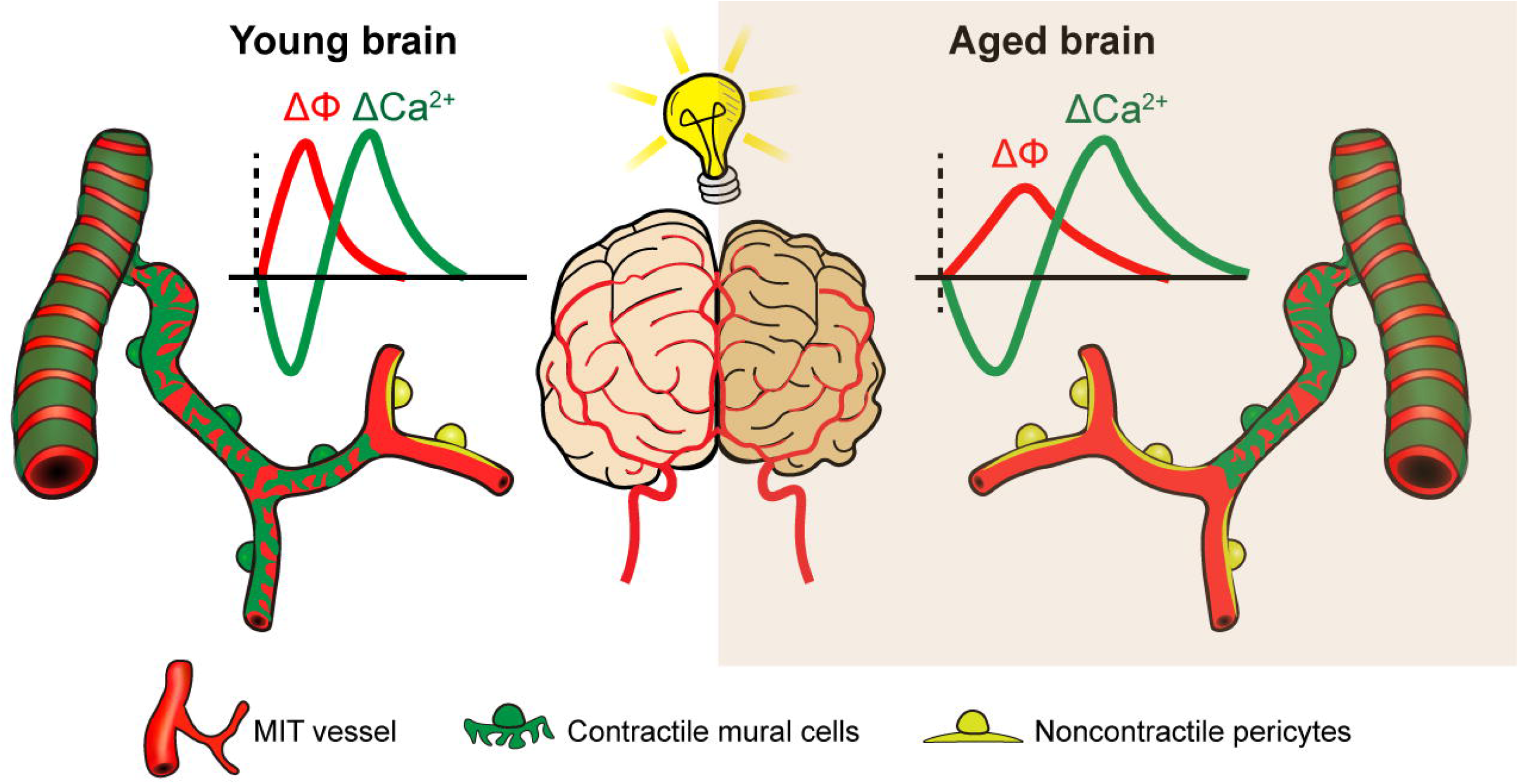
Summary for the remodeled neurovascular coupling function in awake aged brain. In the awake aged brain, vascular mural cells exhibit reduced calcium sensitivity, decreased handling speed of calcium ions, and retracted transition point from contractile to non-contractile mural cells at the capillaries. These factors collectively result in a remodeled neurovascular coupling in response to neuronal stimulus, which shifting from the “fast and strong” mode in young brains to the “slow but prolonged” mode in aged brains.

Studies on the effects of aging on neurovascular coupling (NVC) have shown varying outcomes in both humans and laboratory animals. Some studies reported reduced effects ^34–37^, others showed no change ^10, 11, 38^, and some studies suggested increased NVC responses as compared with young adults ^39, 40^. This variability could be attributed to differences in experimental methods, age of animals, and the specific brain states. In our study, we used awake animal imaging techniques with mice above 20 months old which is equivalent to >60 years in human age ^41^. In the old awake mice, we found that the NVC response displayed a remodeled rather than impaired pattern: from ‘strong and fast’ mode in young brains to ‘slow but prolonged’ mode in aged brains. Under this strategy, the functional hyperemia elicited by a single stimulus in old brains was preserved. This maintains the overall integrity of sensory functions in the aging brain and is consistent with the compensatory scaffolding theory, according to which aged brain compensates for adverse effects of aging to maximize the preservation of brain function ^42^.

The MIT that comprises penetrating arterioles and pre-arteriolar capillaries is thought to play a central role in NVC. Especially the initial branches of capillaries, which are covered by contractile pericytes, are crucial in maintaining capillary perfusion ^43–47^. Our previous study using anesthetized mice showed age-related reduction in vascular reactivity that was most pronounced at precapillary sphincters and first few branches of capillaries ^4^. Our current results indicated that in awake old mice, the vasodilation was more pronounced at the initial capillary segments as compared with penetrating arterioles, indicating that the initial capillaries are the first to be affected by aging. Capillaries account for ∼90% of total cerebral blood vessels, and most of them can only passively dilate due to lack of αSMA-expresseing mural cell coverage ^48, 49^. Thus, the remodeled vasodilation pattern occurring at the initial segment of capillaries is crucial for influencing the perfusion pattern in the entire downstream capillary network, optimally compensating for the energy demands of neurons in aged brains. However, under high-intensity stimulation (spontaneous locomotion accompanied with whisking), the ‘slow but prolonged’ vasodilation pattern may not provide sufficient energy supply that is needed.

In awake animals at resting state, VMCs maintain a relatively high [Ca^2+^]_i_ compared with other brain cells to keep a tonic contraction, contributing to the maintenance of vascular basal tone and regulation of blood flow ^50^. Our results showed a similar resting vessel diameter at MIT in both adult and old mice, which suggested a similar vascular basal tone under resting conditions, while the age-related decrease of Ca^2+^ handling speed may indicate changes in Ca^2+^ transport mechanisms. A previous *in vitro* study, which showed that the overall Ca^2+^ extrusion activity in aged smooth muscle cells decreased, supports this hypothesis ^51^. Our current data show that the amplitude of calcium signal fluctuations in VMCs is similar between the old and adult groups, suggesting the presence of compensatory mechanisms, leading to insignificant changes in the calcium capacity of VMCs during aging. Besides, calcium-independent mechanisms associated with altered calcium sensitization may also modulate vessel activities during aging. Decreased calcium sensitivity indicates that there may be calcium independent signaling involved in the regulation of SMC contractility in aged mice. The reduced activity of PKA/PKG in aging may be one of the reasons for the decreased relaxation of aged SMCs in response to stimuli ^52–54^. However, the mechanisms leading to the prolonged duration of vasodilation still need to be further explored.

Anesthetics have been widely used in the studies of neurovascular coupling in live animal models. However, experiments using anesthetized animal models introduce difficulties in the interpretation of neurovascular coupling mechanisms, as various anesthetics target different action sites ^55, 56^. Besides, anesthesia also leads to changes in the synaptic transmission of the cortical neuronal circuit ^57–60^, as well as brain resting metabolism and neurovascular coupling ^13, 61,62^. Therefore, conclusions drawn from experiments with un-anesthetized and naturally behaving animal models are more closely aligned with the natural physiological conditions. In the experiments with awake mice, we report that the decline in vascular responsivity in old mice is primarily manifested as a reduced dilation speed, and that this decline was much milder in awake mice as compared to results obtained under anesthesia. Furthermore, although previous studies have shown that vascular tone may be impaired in old brains and lead to a large basal vessel diameter ^4, 63, 64^, we found that the resting vessel diameter in awake old mice remained unchanged as compared to adult mice. This suggests that in the awake state, the vascular tone in aged brain is maintained.

Another advantage of conducting experiments with awake mice is the simultaneous recording and observation of mouse behavior. Mouse pupil size dynamics are often used as readout of the level of arousal in mice, which strongly affects sensory processing and behavior ^65–67^. In our experiments, we observed similar amplitude and duration of pupil dilation caused by the whisker air-puff stimulation in adult and old mice, suggesting that somatosensory input is unimpaired during aging. This was further confirmed by monitoring whisking movement induced by air-puff stimuli. In all animals, another noticeable behavior change during aging is the deterioration of motor abilities ^68^. We observed a significant decline in the spontaneous locomotion of old mice and a corresponding reduction in vascular response. As compared to whisking elicited by air-puff stimulation, spontaneous locomotion is an endogenous activity involving multiple brain regions ^69^, which is expected to elicit stronger NVC responses as compared whisking alone. This was evident in the prolonged duration of whisker movements induced by locomotion compared with whisker air-puff. In this scenario, the NVC in the aging brain was incapable of producing a similar response as that in the young brain, indicating limitations in the compensatory strategy. Taken together, our data obtained in awake animals suggest the remodeling of neurovascular coupling in the aging brain, which is reflected in the calcium dynamics of aged VMCs. Our data provide new insights to adaptive strategies for the aged brain, which may be useful for strategies that aim to mitigate age-related cerebrovascular pathologies.

## Acknowledgments

This study was supported by the Lundbeck Foundation, the Independent Research Fund Denmark, the Novo Nordisk Foundation, and a Nordea Foundation Grant to the Center for Healthy Aging. We thank Lars Jørn Jensen for scientific discussion. We thank Agnete Kirkeby, Tune Pers for constructive feedback and valuable suggestions on improving the experiments. We thank the technical assistance of Core Facility for Integrated Microscopy, University of Copenhagen and Flow Cytometry & Single Cell Core Facility, University of Copenhagen and Single-Cell Omics Platform of Center for Basic Metabolic Research, University of Copenhagen. We thank the Shanghai Jiao Tong University for the scholarship support.

## Author Contributions

C.C. and M.L. designed the research and directed the work; X.Z. performed most of the experiments, including chronic cranial window surgery, laser speckle contrast imaging, two-photon imaging, immunohistochemistry, cells purification, pupillometry and whisking detection. L.T., C.C. and C.H. performed two-photon imaging. A.H.N., C.M.G. and X.H. performed quantitative PCR analysis, immunohistochemistry and pupillometry. Y.D., D.P., and I.A. contributed analytic tools for laser speckle contrast imaging and computational modeling; X.Z., C.C. and Y.D. analyzed data; X.Z., C.C. and M.L. wrote the paper. All of the authors revised the final version of manuscript.

## Supplemental material

Supplemental material for this article is available online.

