## Supplemental information for "The neurovascular coupling response of the aged brain is brain-state dependent"

**Supplementary figures**

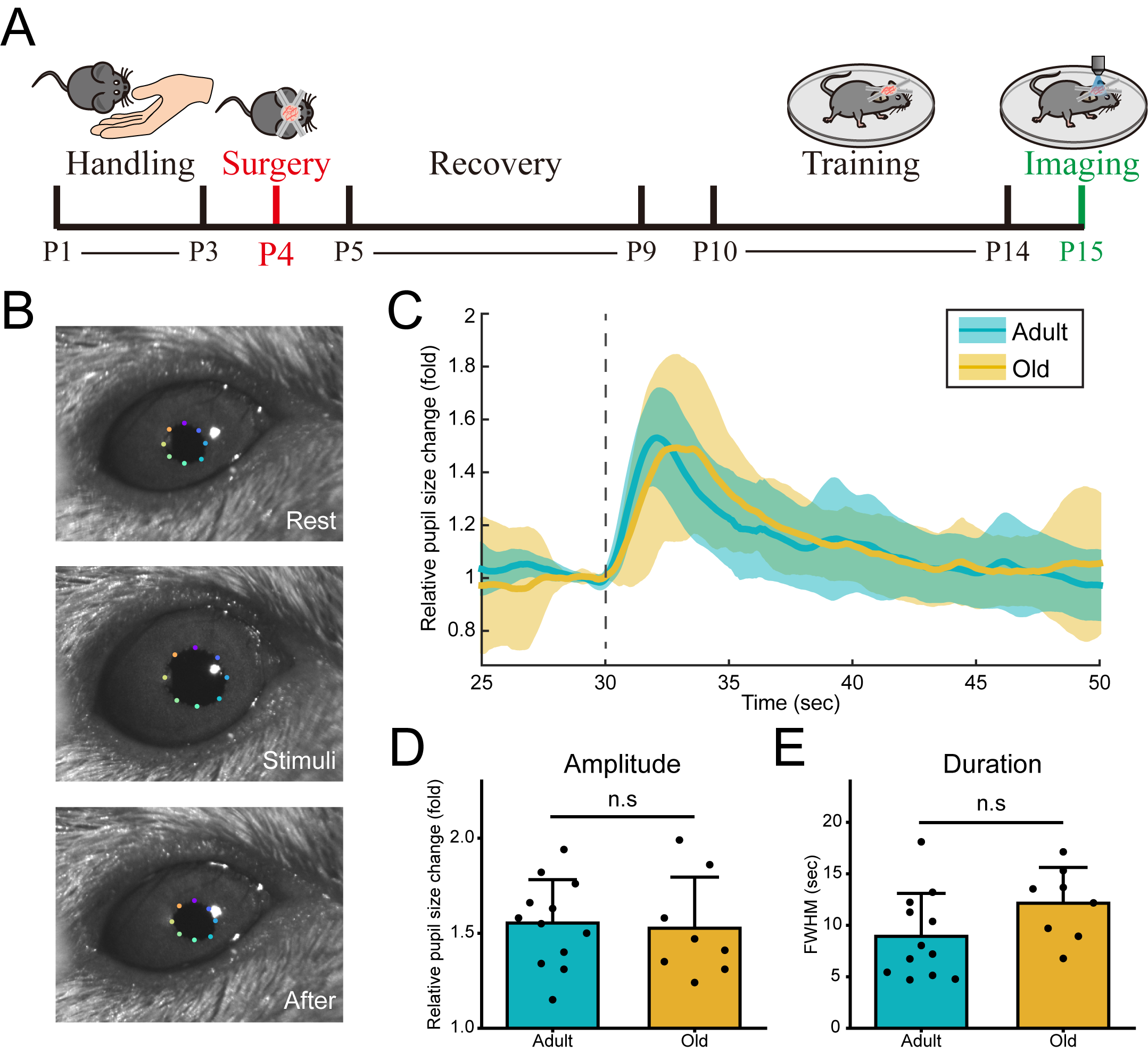

Figure S1. The similar sensory activation in adult and aged mice.

(A). Diagram of the awake mouse imaging preparation process. Chronic cranial window surgery was performed on mice after three days of familiarization with the environment during handling. After the surgery, mice were given a rest period of 5 days and receive postoperative care. Subsequently, continuous adaptive training was conducted for another 5 days and followed by imaging experiments.

(B). Representative images of mouse pupillometry upon air-puff stimulation. Note that the eight dots indicate the edge positions of the mouse pupils.

(C). Mean (solid curve) and SD (shadow) traces of the pupil size change upon air-puff stimulation in adult and old mice. The dashed line indicates the time when the air-puff was delivered.

(D and E). Comparison of relative pupil dilation amplitude (D) and duration (E) in adult and old mice. N = 12 recordings from 4 adult mice and 8 recordings from 3 old mice. Data are presented as mean values ± SD. In (D and E), Unpaired t test. n.s, not significant.

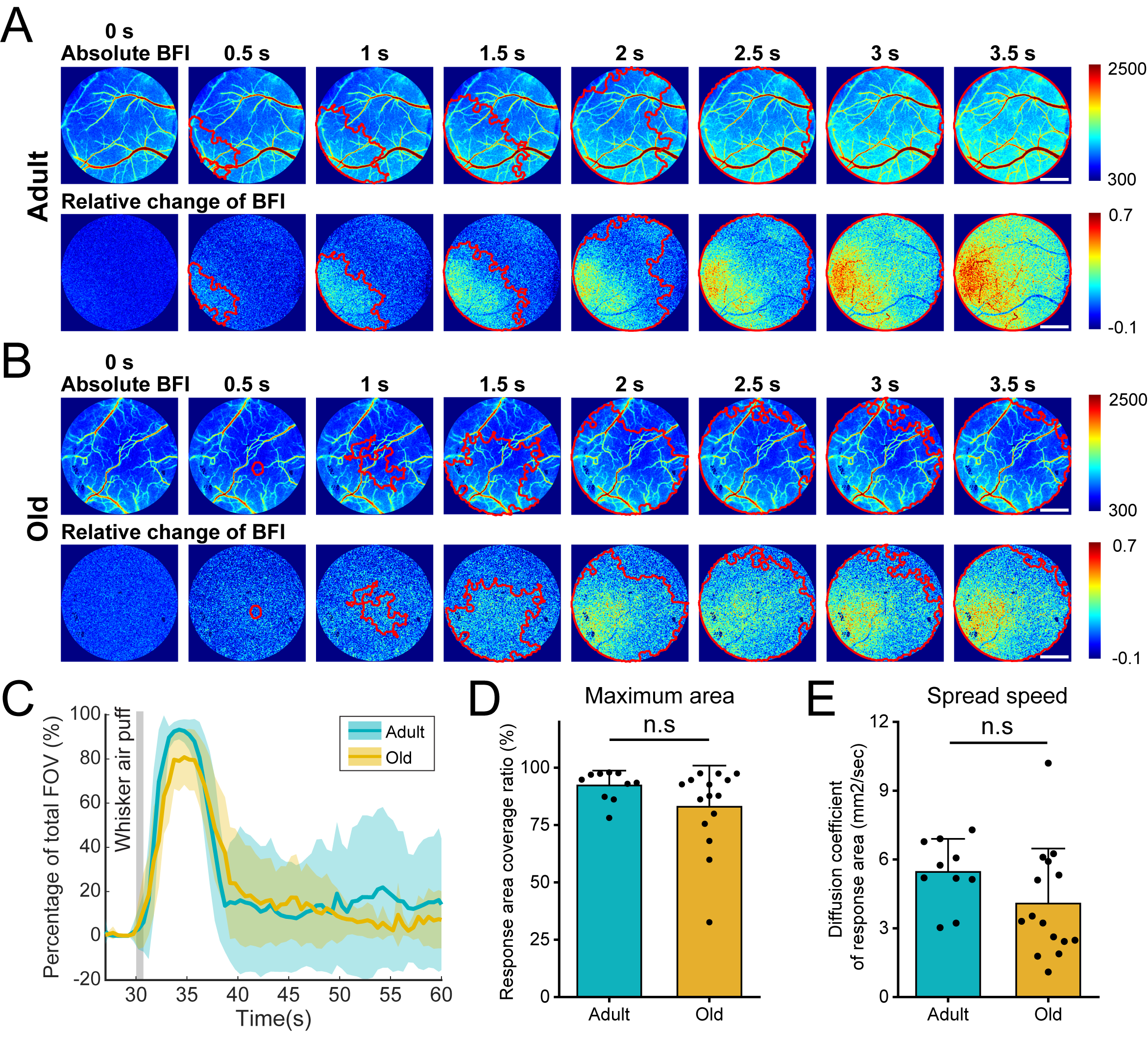

Figure S2. No difference in response zone area and spread speed between adult and aged mice under LSCI.

(A and B). Representative images of cortical blood flow change in response to air-puff stimulation to show the spread of the response zone in adult and old mice. The upper row shows real-time absolute blood flow index, the lower row shows real-time per-pixel relative blood flow changes. The response zone area is indicated by red ROIs. Scale bar, 1 mm.

(C). Mean (solid curve) and SD (shadow) traces show the temporal changes in the percentage of responsive zone to the total FOV area following air puff stimulation in adult and old mice.

(D-E). Bar graphs showing the ratio of maximum responsive zone area to the total FOV area (D) and the diffusion coefficient of response zone (E) in response to air-puff stimulation in adult and old mice. N= 10 recordings from 7 adult mice and 15 recordings from 7 old mice. Data are presented as mean values ± SD. In (D), Mann-Whitney test; in (E), Unpaired t test. n.s, not significant.

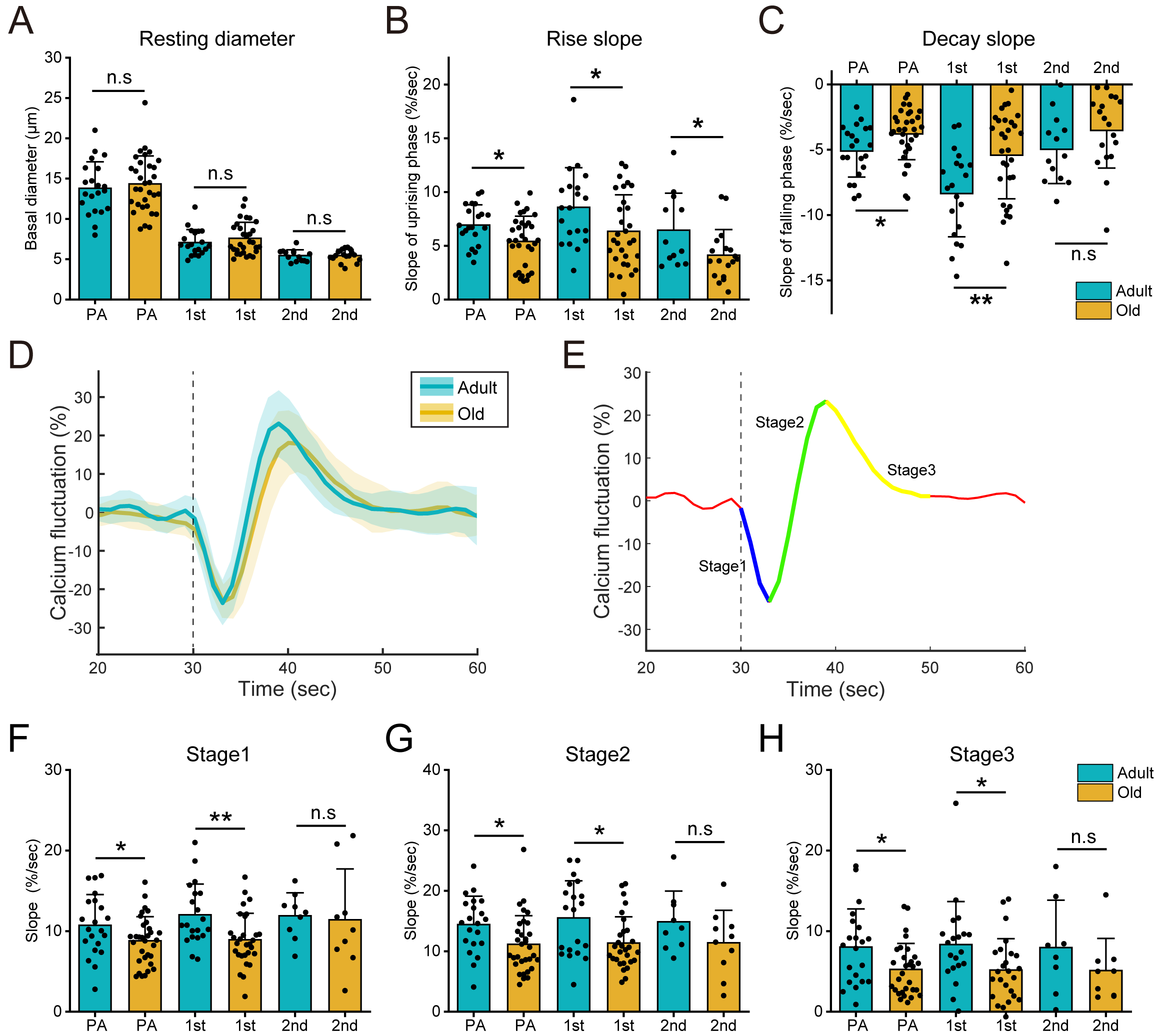

Figure S3. Vasodilation velocity and vascular mural cell calcium transport speed decreases in aged mice.

(A). Bar graphs showing the basal diameter for each vessel order at resting state in adult and old mice. N = 21 vessels from 8 adult mice and 32 vessels from 7 old mice. Data are presented as mean values ± SD. Unpaired t test was used. n.s, not significant.

(B and C). Comparison of rise (10%-90%) and decay (90%-37%) slope of vessel dilation curve at each location in adult and old mice. N = 21 vessels from 8 adult mice and 32 vessels from 7 old mice. Data are presented as mean values ± SD. Unpaired t test was used. *p < 0.05; **p < 0.01; n.s, not significant.

(D and E). Representative curves showing the calcium signal dynamics in vascular mural cells upon air-puff stimulation in adult and old mice. The calcium signal fluctuations are divided into three stages based on oscillations. Stage 1: from baseline to negative peak; Stage 2: from negative peak to positive peak; Stage 3: from positive peak back to normal.

(F-H). Comparison of the slopes of vascular mural cell calcium fluctuation in three stages in adult and old mice. Note that the slope values represent the 10%-90% slope in stage 1 and 2, and the 90%-37% slope in stage 3. N = 21 vessels from 8 adult mice and 32 vessels from 7 old mice. Data are presented as mean values ± SD. In (F, G_PA_ and H_PA_), Unpaired t test; In (G_2nd_ and H_2nd_), Unpaired t test with Welch correction; in (G_1st_ and H_1st_), Mann-Whitney test. *p < 0.05; **p < 0.01; n.s, not significant.

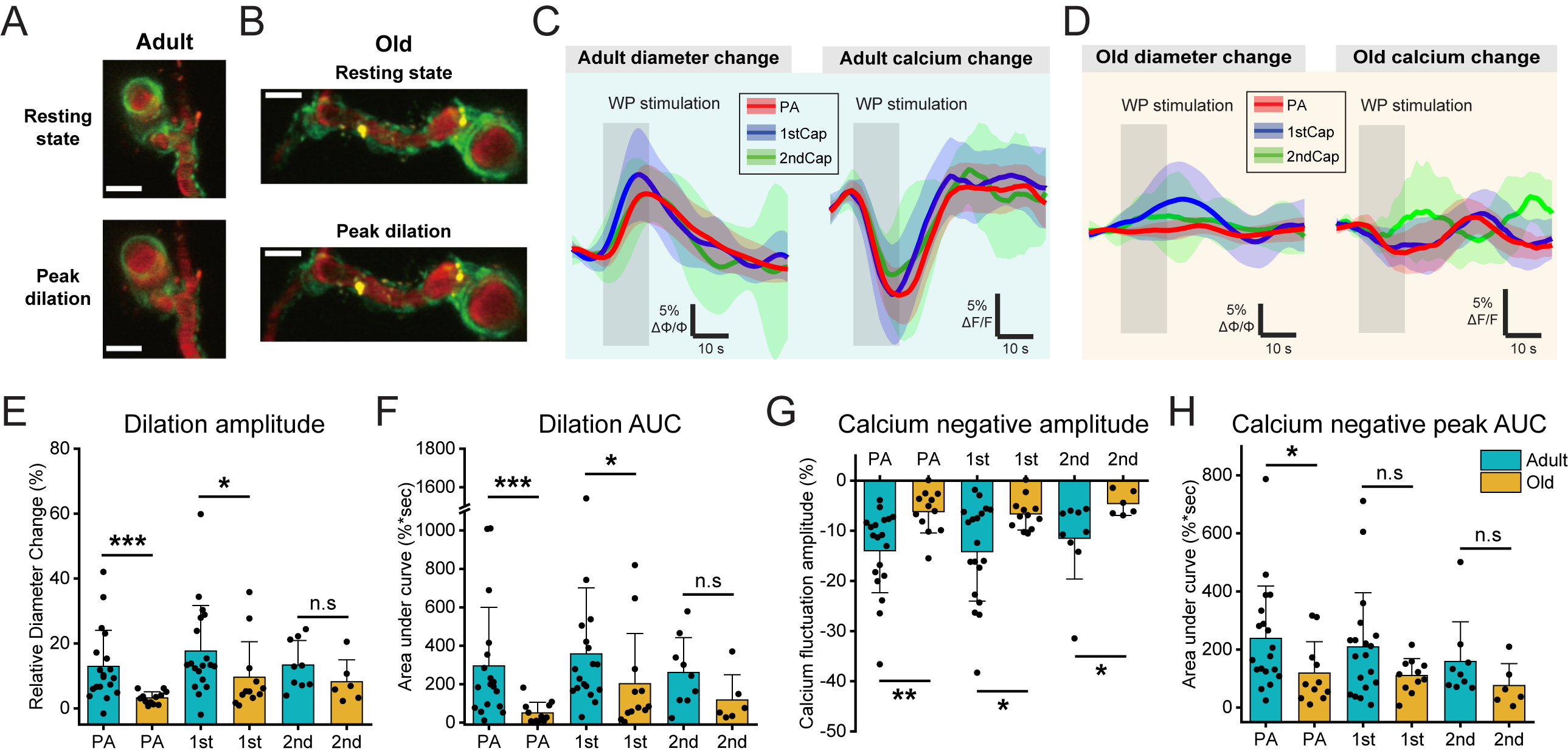

Figure S4. Neurovascular coupling function decreases in aged anesthetized mice.

(A and B). Representative images showing the response of blood vessels in MIT to whisker-pad stimulation in adult and old anesthetized mice. Scale bar: 10 μm. Note that the vessel lumen was labelled with Texas Red dextran, and the vascular mural cell expressed GCaMP8.1.

(C and D). Mean (solid curve) and SD (shadow) traces of the vessel diameter change (C) and calcium fluctuation in vascular mural cells (D) at each vessel location during whisker-pad stimulation in adult and old anesthetized mice.

(E and F). Bar graphs showing the amplitude and area under the curve of vessel diameter change induced by whisker-pad stimulation in adult and old anesthetized mice. N = 19 vessels from 3 adult mice and 12 vessels from 3 old mice. Data are presented as mean values ± SD.

(G and H). Bar graphs summarizing the negative amplitude and area under the curve of mural cell calcium response induced by whisker-pad stimulation in adult and old anesthetized mice. N = 19 vessels from 3 adult mice and 12 vessels from 3 old mice. Data are presented as mean values ± SD.

In (E_PA & 1st_, F_PA & 1st_, G_PA & 1st_ and H), Mann-Whitney test; In (E_2nd_ and F_2nd_), Unpaired t test; in (G_2nd_), Unpaired t test with Welch correction. *p < 0.05; **p < 0.01; ***p < 0.001; n.s, not significant.

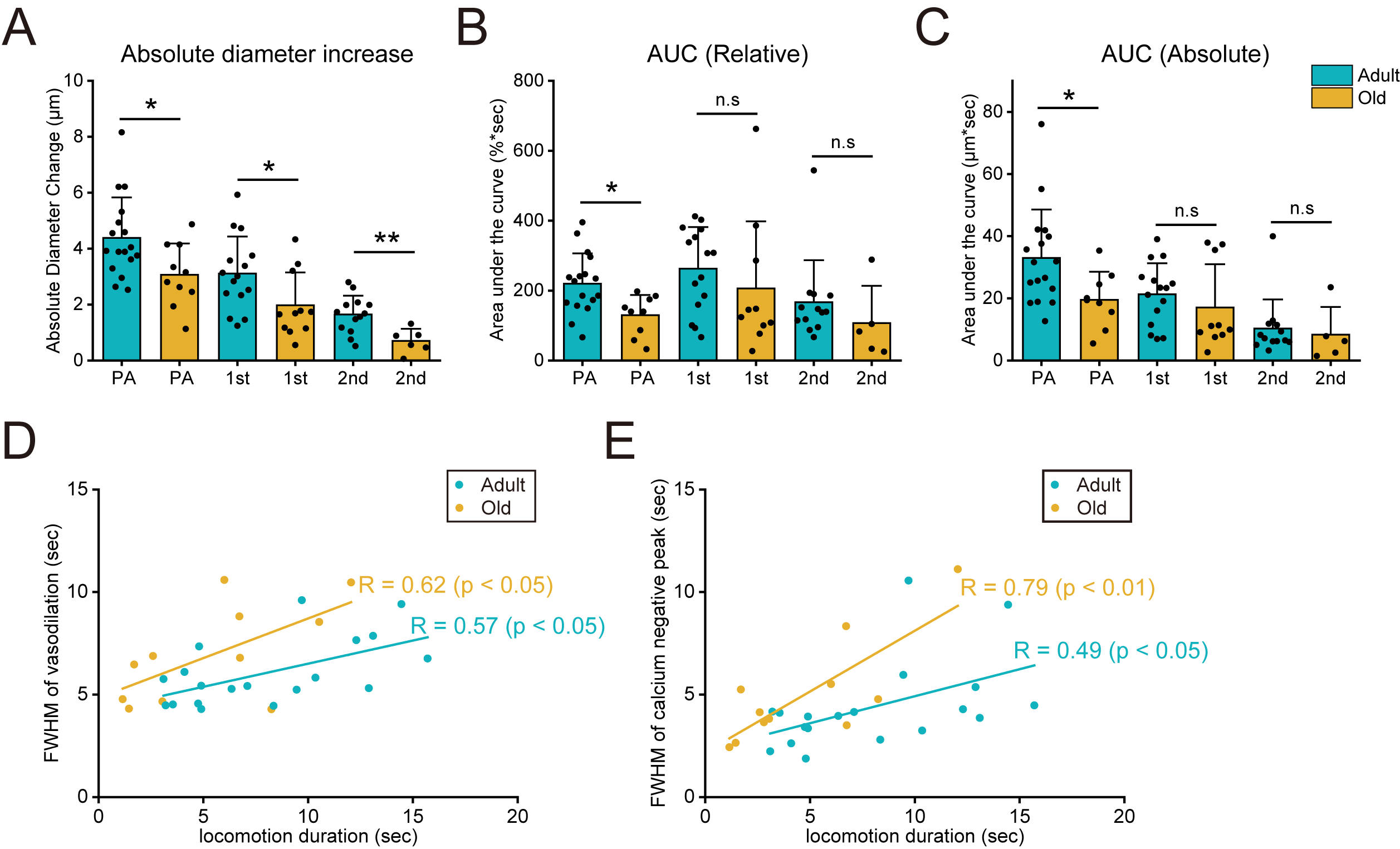

Figure S5. Locomotion induced neurovascular coupling response is reduced in aged mice.

(A). Bar graph showing the absolute diameter change amplitude at each vessel location during spontaneous locomotion in adult and old mice. N = 19 recordings from 6 adult mice and 11 recordings from 4 old mice. Data are presented as mean values ± SD. In (A_PA & 2nd_), Unpaired t test; in (A_1st_), Mann-Whitney test. *p < 0.05; **p < 0.01.

(B and C) Bar graphs summarizing the relative and absolute area under the curve of spontaneous locomotion induced vessel dilation at each location in adult and old mice. N = 19 recordings from 6 adult mice and 10 recordings from 4 old mice. Data are presented as mean values ± SD. In (B_PA_ and C_PA_), Unpaired t test; in (B_1st & 2nd_ and C_1st & 2nd_), Mann-Whitney test. *p < 0.05; n.s, not significant.

(D and E). The correlation analysis of the relationship between spontaneous locomotion duration with vasodilation duration (D) and mural cell calcium negative peak duration (E) of vasculature in adult and old mice. The correlation analysis is presented using linear fitting and Pearson’s r value is indicated. N = 19 recordings from 6 adult mice and 11 recordings from 4 old mice.

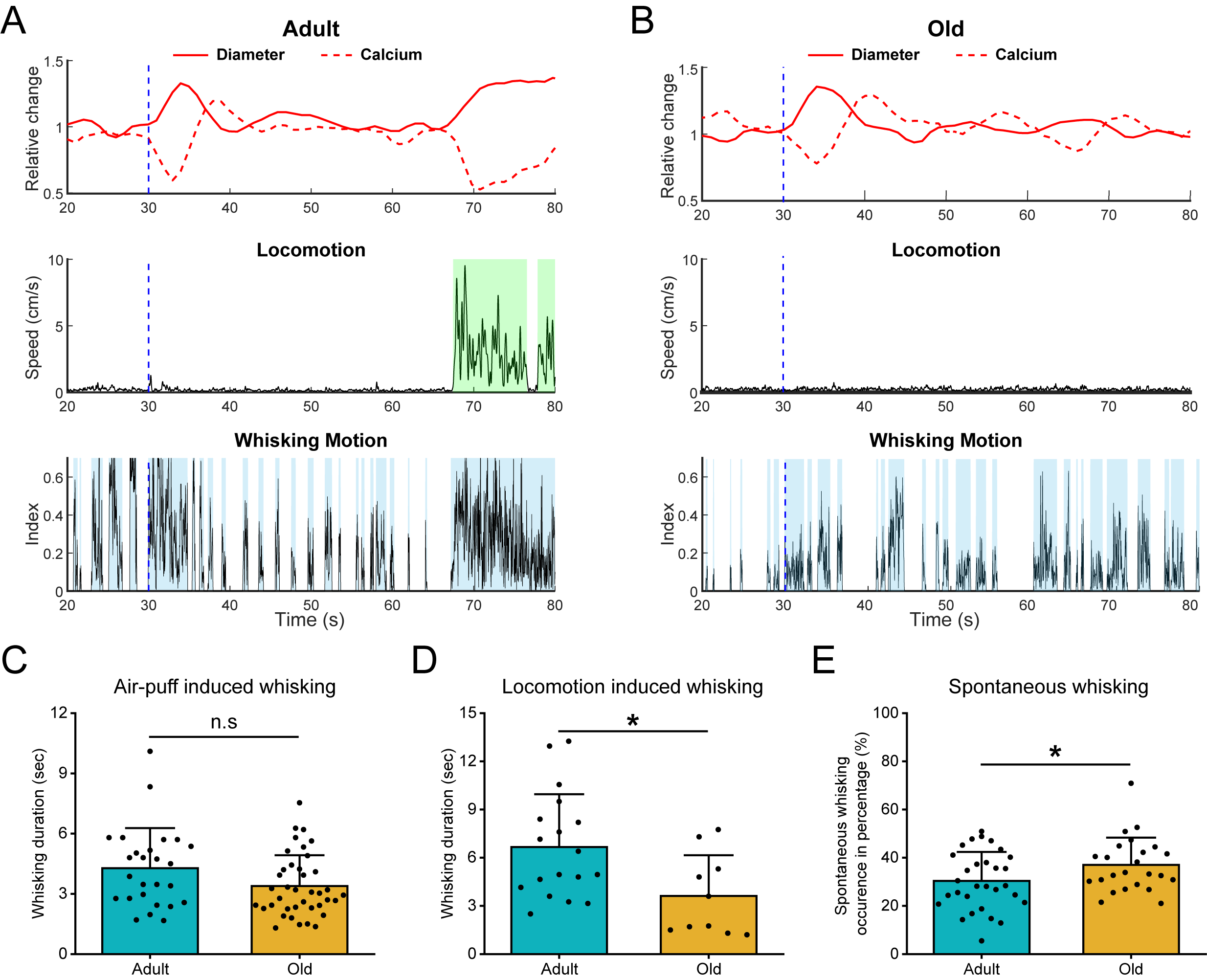

Figure S6. Evoked and spontaneous whisker movement detection in adult and aged mice.

(A and B). Representative traces showing the whisking detection (blue shadow) during the resting state, air-puff stimulation (blue dashed line) and spontaneous locomotion (green shadow) in adult and old mice.

(C-E). Comparison of whisking occurrence upon air-puff stimulation (C, n = 26 recordings in adult mice and 39 recordings in old mice), induced by spontaneous locomotion (D, n = 19 recordings in adult mice and 11 recordings in old mice) and during resting state (E, n = 30 recordings in adult mice and 24 recordings in old mice) in adult and old mice. Note the whisking occurrence during resting state is presented as a percentage due to the variations in the duration of the resting state in different mice. Data are presented as mean values ± SD. In (C and D), Mann-Whitney test; in (E), Unpaired t test. *p < 0.05; n.s, not significant.

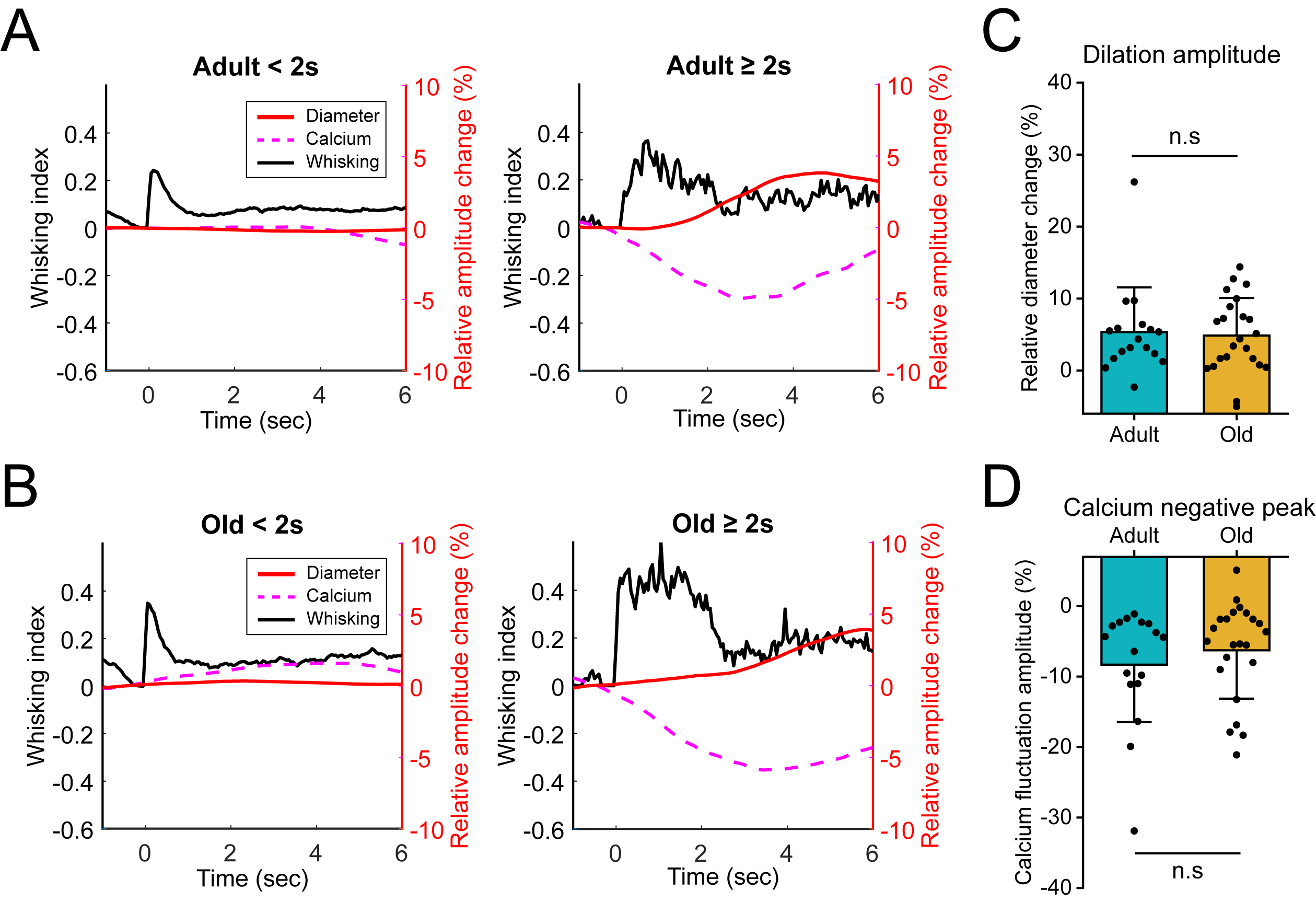

Figure S7. Spontaneous whisking induces same level of neurovascular coupling response in adult and aged mice in resting state.

(A and B). Mean traces of short (< 2 s) or long (≥ 2 s) spontaneous whisking, along with the corresponding diameter changes and mural cell calcium response of vasculature.

(C and D). Bar graphs show the amplitude of vessel dilation (C) and mural cell calcium response (D) induced by long spontaneous whisking (≥ 2 s). N = 17 events in adult mice and 23 events in old mice. Data are presented as mean values ± SD. Mann-Whitney test was performed and ‘n.s’ indicates not significant.

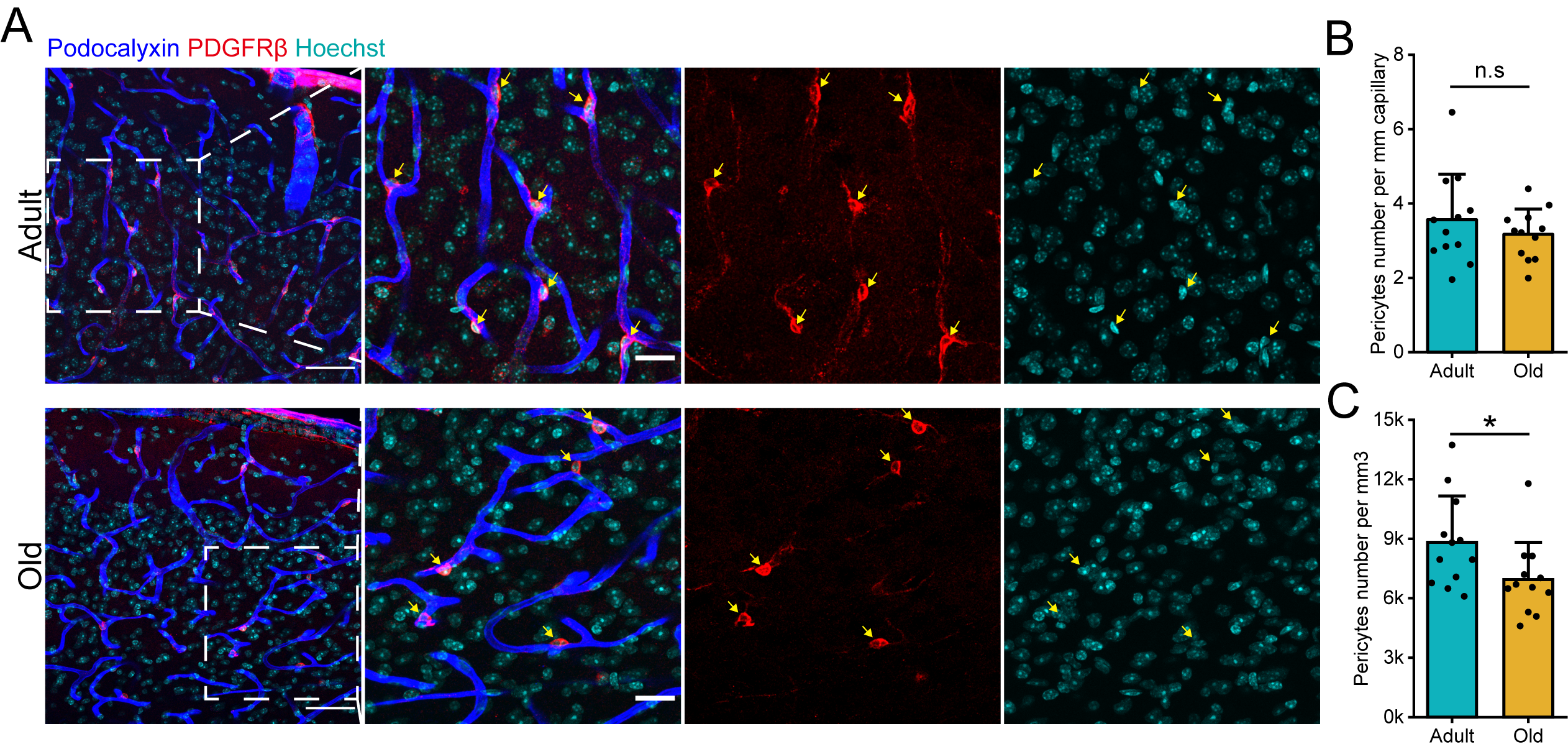

Figure S8. Comparison of mural cell calcium pump gene expression and pericyte density in adult and aged mice.

(A). Immunostaining of pericytes in adult and old mice cortex. The arrows mark pericytes located on capillaries. Scale bar: 50 μm (left) and 10 μm (inlet).

(B and C). Comparison of pericyte density per mm of capillary and spatial density in the adult and old mice cortex. N = 12 slices for both groups. Data were collected from 3 adult and 3 old mice and are presented as mean values ± SD. Unpaired t test was performed. *p < 0.05; n.s, not significant.
